# Conductive microgel annealed scaffolds enhance myogenic potential of myoblastic cells

**DOI:** 10.1101/2023.08.01.551533

**Authors:** Alena Casella, Jeremy Lowen, Nathan Shimamoto, Katherine H. Griffin, Andrea C. Filler, Alyssa Panitch, J. Kent Leach

## Abstract

Bioelectricity is an understudied phenomenon to guide tissue homeostasis and regeneration. Conductive biomaterials may capture native or exogenous bioelectric signaling, but incorporation of conductive moieties is limited by cytotoxicity, poor injectability, or insufficient stimulation. Microgel annealed scaffolds are promising as hydrogel-based materials due to their inherent void space that facilitates cell migration and proliferation better than nanoporous bulk hydrogels. We generated conductive microgels from poly(ethylene) glycol and poly(3,4-ethylenedioxythiophene) polystyrene sulfonate (PEDOT:PSS) to explore the interplay of void volume and conductivity on myogenic differentiation. PEDOT:PSS increased microgel conductivity over 2-fold while maintaining stiffness, annealing strength, and viability of associated myoblastic cells. C2C12 myoblasts exhibited increases in the late-stage differentiation marker myosin heavy chain as a function of both porosity and conductivity. Myogenin, an earlier marker, was influenced only by porosity. Human skeletal muscle derived cells exhibited increased *Myod1*, IGF-1, and IGFBP-2 at earlier timepoints on conductive microgel scaffolds compared to non-conductive scaffolds. They also secreted higher levels of VEGF at early timepoints and expressed factors that led to macrophage polarization patterns observed during muscle repair. These data indicate that conductivity aids myogenic differentiation of myogenic cell lines and primary cells, motivating the need for future translational studies to promote muscle repair.

## 1. Introduction

Muscle tissue engineering is a promising strategy for repairing large muscle wounds such as volumetric muscle loss (VML) that surpass the body’s innate healing ability. VML and other musculoskeletal disorders, which affect over 500 million people worldwide, may result in reduced mobility, disability, and significant economic burden.^[1,2]^ Autologous muscle graft is the current gold standard of treatment, which has negative side effects of donor site morbidity and atrophy. Muscle tissue engineering provides alternative strategies for healing compared to native muscle grafts.^[3,4]^

Synthetic and natural polymers have been developed for specific applications in muscle tissue engineering including aligned structures to recapitulate muscle isotropy, elastic materials to mimic the contractile function of muscle tissue, and hydrogels for use as volume fillers and cell delivery vehicles. Hydrogels are popular for cell and drug delivery due to their tunability and biophysical behavior that can mimic that of native tissues.^[5]^ Bulk hydrogels have been widely used in muscle tissue engineering due to their ease of handling, cell-friendly nature, and tunable mechanical properties. However, bulk hydrogels are typically nanoporous in nature, which limits cell-cell interaction and cell migration until the surrounding extracellular matrix (ECM) is degraded. Microgels are a promising hydrogel platform due to their modularity and microporosity.^[6]^ Unlike conventional nanoporous bulk hydrogels, the inherent void space between microgels permits immediate cell migration without the need to remodel the local environment. A multitude of studies have examined bulk hydrogels for muscle tissue engineering,^[5]^ but there is limited evidence of exploring microgels for this application. In one example, microgels were mixed with silver nanoparticles to form a conductive mixture that conferred electric signals across *ex vivo* tissues, yet their influence on muscle cell function was not reported.^[7]^

Bioelectricity is a potent but understudied stimulus that plays a key role in muscle tissue formation and function.^[8]^ Biomaterials that possess bioelectric potential (*i.e.*, conductivity) are an exciting strategy to advance the field of tissue engineering. Electrically conductive biomaterials continue to gain popularity owing to their ability to direct cell differentiation and maturation, particularly for nerve^[9]^ and cardiac tissue repair.^[10]^ Synthetic conductive polymers such as polypyrrole, polyaniline, and PEDOT:PSS (poly(3,4-ethylenedioxythiophene) polystyrene sulfonate) or carbon-based materials (*e.g.*, carbon nanotubes, graphene, etc.) are frequently used to imbue hydrogels with electroactive properties.^[8,11]^ For example, synthetic electrospun fibers developed for muscle tissue engineering that contained either polyaniline blends^[12,13]^ or PEDOT:PSS nanoparticles^[14]^ possessed conductivity and improved muscle cell differentiation and maturation. PEDOT was also incorporated into directionally aligned collagen scaffolds to instruct myoblast behavior.^[15]^ While these reports include both electrical and physical cues to promote cell differentiation toward myogenesis, the interplay between conductivity and scaffold porosity has yet to be directly interrogated.

Herein, we aim to combine the microporosity of microgel annealed scaffolds with the conductivity of PEDOT:PSS to enhance myogenic differentiation. We demonstrate a conductive microgel platform which outperforms both conductive bulk degradable and non-conductive microporous scaffolds in promoting myogenic differentiation. We generate microgel scaffolds possessing attributes characteristic of native muscle tissue. Cell viability and scaffold stiffness is not altered by the addition of PEDOT:PSS, and conductive microgels can be annealed into a contiguous scaffold. Gene and protein expression indicative of myogenic differentiation is upregulated in our conductive microgel scaffolds when seeded with murine C2C12 myoblasts or human skeletal muscle derived cells compared to matched control gels, suggesting the importance of both electroactivity and microporosity for muscle regeneration.

## 2. Materials and Methods

### 2.1 Microgel synthesis

Microgels were fabricated using a previously described microfluidic device that was adapted by our group.^[16,17]^ For the production of nondegradable microgels, the aqueous phase consisted of 10 kDa 8-arm PEG-vinyl sulfone (PEG-VS) (JenKem, Plano, TX) and RGD (Ac-RGDSPGERCG-NH_2_, Genscript, Piscataway, NJ) in 100 mM HEPES buffer (N-2-hydroxyethylpiperazine-N’-2-ethanesulfonic acid, pH 5.25, Sigma, St. Louis, MO) mixed with 3.5 kDa PEG-DT (JenKem) dissolved in diH_2_O with or without PEDOT:PSS (PH1000, Ossila, Sheffield, UK). The final microgel concentrations were 4.5 mM PEG-VS, 10.8 mM PEG-DT, 1 mM RGD, and 0.25 wt% PEDOT:PSS. The oil phase consisted of Novec 7500 Oil and 0.75 wt% Picosurf (Sphere Fluidics, Cambridge, UK). After exiting the device, microgels were combined with a solution of 1 v/v% triethylamine (TEA, Sigma) in Novec 7500 Oil using a Y-junction (IDEX Health and Science, Oak Harbor, WA) and left at room temperature overnight to ensure complete crosslinking. Microgels with final diameter of ∼150 µm were cleaned to remove residual oil and surfactant as previously described.^[17]^

### 2.2 Annealing microgels

Microgels were annealed as previously described.^[17]^ Briefly, microgels were suspended in an annealing solution of additional crosslinker in HEPES with 0.4% VA-086 photoinitiator (FUJIFULM Wako Chemicals, Richmond, VA) equal to the aggregate volume of microgels. After incubating for 1 min, the microgels were spun down for 3 min at 14,000 × *g*. The supernatant was removed and microgels were optionally mixed with cells before plating in a 8 mm x 1.5 mm cylindrical mold. The microgel slurry was then exposed to UV light (20 mW/cm^2^, 320-500 nm, Omnicure S2000) for 2 min to form annealed scaffolds.

### 2.3 Bulk degradable gel synthesis

GPQ-A (GCRDGPQGIAGQDRCG, Genscript), a protease-cleavable crosslinking peptide, was substituted for PEG-DT to permit matrix metalloproteinase (MMP)-mediated degradation. The final concentrations were 8 mM PEG-VS, 19.2 mM GPQ-A, 1 mM RGD, and 0.25 wt% PEDOT:PSS. A precursor solution consisting of PEG-VS, RGD, and optionally PEDOT:PSS in HEPES (25 mM, pH 7.2) at 2X concentration was combined with cells and pipetted into the desired mold. An equal volume of 2X GPQ-A (pH 8.3) in media was then mixed in by pipetting up and down. The gels were incubated at 37°C for 15 min before being transferred to a well plate.

### 2.4 Conductivity testing

Annealed microgels were electrically characterized as described previously.^[18]^ Briefly, 8 mm scaffolds were constrained by a PDMS mold and sandwiched between two brass plates. The sandwich was then stabilized between the jaws of a tabletop angle vise using PDMS blocks as a barrier between the plate and the jaw. One brass plate was connected to a power supply (BK Precision 1735A, Yorba Linda, CA) using alligator clips and the other plate was connected to a multimeter (SparkFun Electronics, Niwot, CO) to measure output current. Voltages ranging from 100 to 500 mV, chosen to avoid the electrolysis of water, were applied to obtain current-voltage curves. After testing, hydrogel diameter and thickness were measured with calipers and hydrogel cross-sectional area was calculated. Current-voltage curves were analyzed for linearity and datasets with an R^2^ value ≥0.9 were accepted for resistance calculations. Conductivity was calculated using Pouillet’s law (**Equation 5.1**). Hydrogels for conductivity testing were stored in ultrapure water to eliminate the confounding effects of ions in other solutions.

#### Equation 5.1

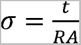

where σ is conductivity in S/cm, t is thickness of the hydrogel (cm), R is resistance (Ω), and A is cross-sectional area (cm^2^).

### 2.5 Mechanical testing

#### 2.5.1 Determination of bulk hydrogel shear storage modulus

Bulk hydrogel scaffolds were loaded onto a Discovery HR2 Rheometer (TA Instruments, New Castle, DE) with a stainless steel, cross hatched, 8 mm plate geometry. An oscillatory strain sweep ranging from 0.004% to 4% strain was performed on each gel using an initial 0.3 N axial force to obtain the linear viscoelastic region (LVR) before failure.

#### 2.5.2 Determination of microgel compressive elastic modulus

Individual microgels were examined using a MicroTester (CellScale, Waterloo ON, Canada). Microgels were loaded onto an anvil in a water bath filled with PBS. The microgels were then compressed to half their diameter by a stainless-steel platen attached to a tungsten rod over 30s. Displacement and force were traced *via* MicroTester software. The slope of the linear region of the compressive modulus *versus* nominal strain graph was recorded as the calculated modulus.^[17,19]^

### 2.6 Cell culture

#### 2.6.1 C2C12 myoblasts

C2C12 murine myoblasts (CRL-1772, Lot #70013341, ATCC, Manassas, VA) were cultured in DMEM (Thermo Fisher Scientific, Waltham, MA) supplemented with 10% FBS (GenClone, San Diego, CA) and 1% penicillin-streptomycin (P/S, Gemini Bio Products, West Sacramento, CA) in standard culture conditions (*i.e.*, 37°C, 5% CO_2_). Cultures were maintained until <70% confluent to prevent myoblast differentiation. Differentiation media was prepared by supplementing DMEM with 2% heat-inactivated horse serum (Thermo Fisher Scientific) and 1% FBS (DMEM-D). Cells were seeded into scaffolds at 1 x 10^6^ cells/mL, and cell-laden scaffolds were cultured in 24-well plates containing growth media for approximately 24 h (0 d) before transferring to a fresh well plate containing differentiation media (1 d). C2C12 proliferation was assessed by measuring DNA content using the Quant-iT PicoGreen dsDNA Kit (ThermoFisher). Differentiation of C2C12 myoblasts in microgel and bulk degradable scaffolds was assessed at 3 and 7 days.

#### 2.6.2 Human skeletal muscle derived cells

Primary human skeletal muscle derived cells (skMDCs) and all associated cell culture reagents were purchased from Cook Myosite (Pittsburgh, PA). The donor was a 29-year-old Caucasian male with BMI of 29 and no history of smoking or diabetes (SK-1111-P01547-29M). Cells were handled according to the manufacturer’s instructions. Briefly, cells were expanded in MyoTonic™ Basal Medium supplemented with MyoTonic™ Growth Supplement. Cells were seeded into microgel scaffolds at 2 x 10^6^ cells/mL. Annealed scaffolds were maintained in growth medium for 1 d before transferring to MyoTonic™ Differentiation media. Muscle cell differentiation and myotube formation was assessed at 1, 3, and 7 d, and media conditioned by these cells was collected for later use.

### 2.7 Immunostaining

Scaffolds were fixed in 4% paraformaldehyde for 1 h and incubated in blocking buffer composed of 10% goat serum (MP Bio, Santa Ana, CA) and 10 mg/mL Bovine Serum Albumin (BSA, Sigma) for 30 min at room temperature. Constructs were then incubated with myosin heavy chain antibody conjugated with Alexa Fluor 488 (1:50; Santa Cruz Biotechnology, 376157) and myogenin antibody conjugated with Alexa Fluor 680 (1:50, Santa Cruz Biotechnology, 12732). Samples were rinsed with PBS and incubated with DAPI (1:500 in PBS; ThermoFisher) for 10 min. Z-stacks were taken on a confocal microscope (Leica Stellaris 5), and max projections were used to illustrate cell morphology throughout the scaffolds.

### 2.8 qPCR

Total RNA was isolated from cells using TRIzol reagent (ThermoFisher) according to the manufacturer’s instructions. RNA quality and quantity was measured using a Nanodrop One© instrument (ThermoFisher) before reverse transcribing to cDNA with the QuantiTect Reverse Transcription kit (Qiagen, Hilden, Germany). All cDNA samples were diluted with PCR-grade ultrapure water to 12.5 ng/µL prior to qPCR. qPCR was performed using Taq PCR Master Mix kit (Qiagen), TaqMan Gene Expression Assay probes (cat. 4331182, ThermoFisher) and a QuantStudio™ 6 instrument (ThermoFisher). Samples were activated at 94°C for 3 min, followed by 40 cycles of 94°C for 30 s, 60°C for 30 s, and 72°C for 1 min, and underwent a final annealing step at 72°C for 10 min.

C2C12 expression of the myogenic differentiation markers MyoD (*Myod1*, Mm00440387_m1), myogenin (*Myog*, Mm00446194_m1) and myosin heavy chain (*Myh7*, Mm00600555_m1) was interrogated at 3 and 7d. Myogenic differentiation and maturation of skMDCs was assessed *via* expression of the same genes (MyoD (*Myod1*, Hs00159528_m1), myogenin (*Myog*, Hs01072232_m1), myosin heavy chain (*Myh7*, Hs01110632_m1)), as well as *Myh2* (Hs00430042_m1), *Pax7* (Hs00242962_m1), and *Ryr1* (Hs00166991_m1) at 1, 3, and 7 d to capture potential differences at earlier timepoints. All genes were normalized to the housekeeping gene GAPDH (Mm99999915_g1, Hs02786624_g1) to yield ΔCt. Gene expression of C2C12s was further normalized to the 0.00% PEDOT:PSS group at 3 d to calculated ΔΔCt. Expression of skMDCs was normalized to that of cells taken prior to microgel seeding. Fold change was calculated using the 2^-ΔΔCt^ method.

### 2.9 Characterization of skMDC secreted factors

Media conditioned by the skMDCs in microgels was collected on 1, 3, and 7 d and frozen at -80°C until analyzed using a Human Cytokine Array C5 kit (Ray Biotech, San Diego, CA) according to the manufacturer’s instructions. Blots were imaged using an Odyssey® XF Imaging System (LI-COR, Lincoln, NE) and normalized against basal MyoTonic™ Differentiation media to account for myokines present in the media alone. Data were analyzed using ImageJ with the Protein Array Analyzer plugin.^[20]^

### 2.10 Endothelial cell tubulogenesis in response to skMDC-secreted factors

Human microvascular endothelial cells (HMVECs) were cultured using Endothelial Growth Media 2 (EGM-2, PromoCell, Heidelberg Germany) prior to staining with CellTracker^TM^ dye (ThermoFisher) and seeding into Matrigel-coated ibidi wells (ibidi, Fitchburg, Wisconsin). Each well contained 1 x 10^4^ cells and was treated with skMDC-conditioned media (described above) such that the ratio of serum-containing EGM-2 to skMDC-CM was 1:4. After 6 h, three images per well were taken using confocal microscopy (Leica Stellaris 5). Images were analyzed for average branch length and total branch length using the Analyze HUVEC Fluo function of the Angiogenesis Analyzer plugin for ImageJ.^[21]^

### 2.11 Flow cytometry for macrophage polarization in response to skMDC-secreted factors

Human THP-1 monocytes were expanded in RPMI 1640 medium (ATCC Formulation: L-glutamine, HEPES, sodium pyruvate, and high glucose) supplemented with 10% FBS (GenClone) in suspension culture under standard culture conditions until the density reached approximately 1x10^6^ cells/mL. Cells were then seeded at 75% confluency and treated with 320 nM phorbol-12-myristate-13-acetate (PMA) for 36 h to induce adherence and differentiation into macrophages.

Following differentiation, THP-1 macrophages were rinsed three times with PBS, treated with conditioned media from skMDCs in microgels after 1, 3, and 7 d in culture at a 1:1 ratio with basal media, and incubated for another 24 h. Cells were then collected for flow cytometry. Polarization controls were treated the same, but instead of conditioned media treatments, macrophages were treated with basal media (M0), 200 ng/mL LPS (M1), and 20 ng/mL IL-4 (M2) (*data not shown*).

Cells were collected with ice cold 2.5 mM EDTA in PBS and gentle scraping. Cells were spun down and resuspended in 37°C 3% FBS in PBS. Cells were then stained for flow cytometric analysis. Following Fc_*γ*_ receptor blocking (1:40, TruStain FcX, BioLegend), cells were stained with antibodies against CD11b (1:40, eBioscience #47-0118-42), HLA-DR (1:40, eBioscience #48-9956-42), and CD206 (1:33, eBioscience #12-2069-42), and CD163 (1:50, Invitrogen #MA5-17719). Cellular viability was evaluated with fixable Zombie Aqua (1:250, Life Tech). Cells were then fixed with 2% PFA, and analyzed on the flow cytometer (Attune NxT, ThermoFisher). Macrophages with an M1 phenotype were characterized by HLA-DR^+^ populations and M2 phenotype by CD206+CD163+ populations.

### 2.12 Statistical analysis

Data are presented as means ± standard deviation. GraphPad Prism 9 software was used to plot all graphs and perform statistical testing. Statistically significant groups are denoted by two conventions: asterisks were used to denote differences when using a t-test or 1-way ANOVA, whereas letters were used when a 2-way ANOVA was employed. Groups denoted by different letters are statistically different. Interactions between time, porosity, and conductivity on gene expression were further analyzed using MATLAB.

## 3. Results

### 3.1 PEG microgels can be imbued with conductivity and annealed into scaffolds

PEG microgels were fabricated as previously described and covalently modified with RGD to facilitate cell adhesion.^[17]^ PEDOT:PSS was optionally introduced into the aqueous phase of microgel fabrication such that the final concentration within each microgel was 0.25 wt% (**Fig. 1A**). Microgels were successfully annealed into 8 mm scaffolds with good retention of PEDOT:PSS, as depicted in **Figure 1B**. The addition of PEDOT:PSS resulted in a significant increase in annealed scaffold conductivity from 1.58 ± 0.68 x 10^-6^ S/cm to 3.52 ± 0.96 x 10^-6^ S/cm (*p*≤0.01; **Fig. 1C**). Next, we assessed if the addition of PEDOT:PSS affected the mechanical properties of the microgels and annealed scaffolds. Compressive modulus of individual microgels was ∼28 kPa for both groups (**Fig. 1D**). Using Hooke’s law for isotropic materials and a Poisson’s ratio of approximately 0.5, the storage modulus of each microgel is estimated as 10 kPa.^[22]^ Since cells interact directly with individual microgels, these mechanical properties were considered appropriate for muscle tissue engineering applications.^[23]^ The storage modulus of annealed scaffolds was not significantly different between groups, indicating that PEDOT:PSS did not affect annealing ability (**Fig. 1E**). Together, these data illustrate successful incorporation of PEDOT:PSS into PEG microgels without altering mechanical properties. The microgels can be annealed to form scaffolds with decoupled electrical and mechanical properties suitable for muscle tissue engineering.

**Figure 1:**
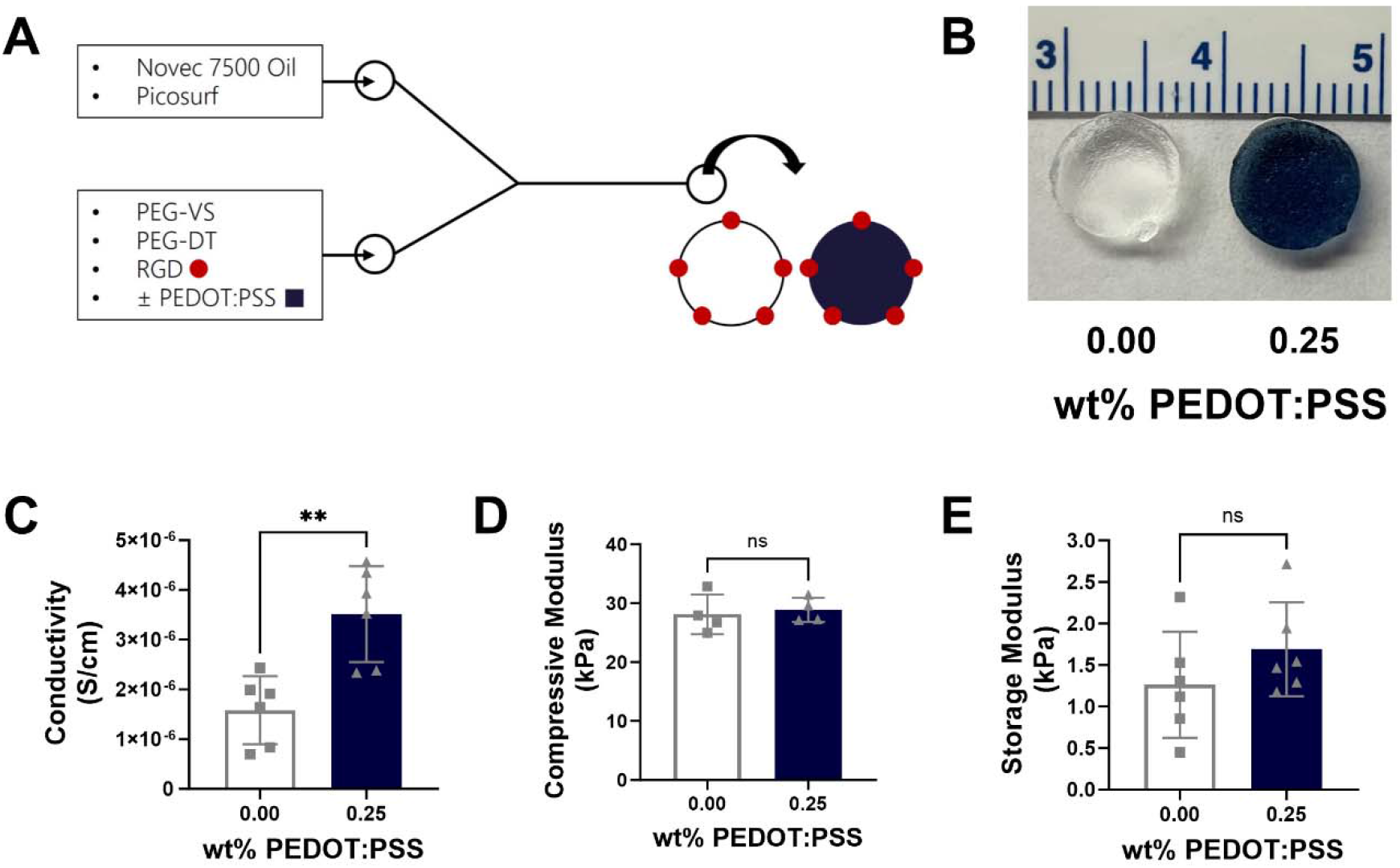
The electrical and mechanical properties of conductive microgels can be decoupled. **(A)** Schematic of PEG microgel modification and PEDOT:PSS incorporation. **(B)** Gross images of 8 mm scaffolds demonstrate successful annealing with UV light and retention of PEDOT:PSS. **(C)** Scaffolds containing PEDOT:PSS were significantly more conductive than non-conductive controls (n=6). PEDOT:PSS did not affect **(D)** the compressive modulus of individual microgels (n=4) or **(E)** the storage modulus of annealed microgel scaffolds (n=6). Groups were compared using a two-tailed t-test where \*\**p*≤0.01 and ns = not significant.

### 3.2 Conductive microgel scaffolds promote C2C12 proliferation

C2C12 mouse myoblasts were seeded into annealed microgel scaffolds or bulk degradable hydrogels as a control. Cells in conductive annealed microgel scaffolds exhibited greater proliferation at 3 days than those in the non-conductive control, evidenced by the increase in DNA content (**Fig. 2A**). There were no changes in DNA content for myoblasts in annealed microgel scaffolds at 7 days.

**Figure 2:**
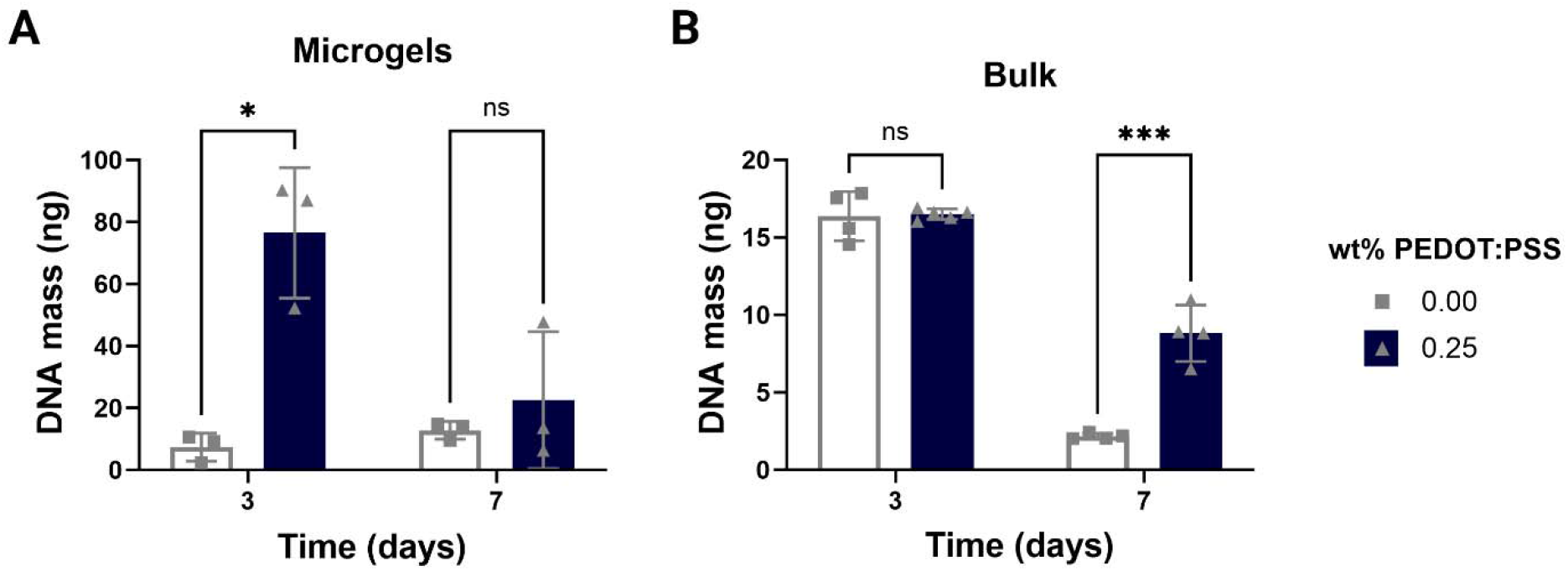
Microgel structure promotes C2C12 proliferation compared to bulk hydrogels. **(A)** DNA content of C2C12s in annealed microgel scaffolds demonstrates conductive scaffolds promoted myoblast proliferation at 3 days (n=3, \**p*≤0.05). **(B)** C2C12s cultured in bulk degradable hydrogels had lower proliferation overall, though the conductive hydrogels supported myoblasts better at 7 days than non-conductive controls (n=4, \*\*\**p*≤0.001). Groups were compared using multiple, unpaired t-tests where ns = not significant.

Conversely, C2C12s grown in bulk degradable gels had no difference in DNA content or metabolic activity at 3 days, regardless of PEDOT:PSS content. At 7 days, however, PEDOT:PSS-containing bulk gels exhibited greater DNA content, indicating that conductive substrates may better support cell viability (**Fig. 2B**). Importantly, DNA content in annealed microgel scaffolds was higher than in bulk hydrogel controls. These data suggest microgels containing PEDOT:PSS support increased cell viability and proliferation of C2C12s compared to microgels without the conductive additive. Furthermore, these data demonstrate the advantage of microgel annealed scaffolds in supporting cell proliferation compared to nanoporous bulk hydrogels that are frequently used for tissue engineering studies.

### 3.3 Microporous structure and conductivity promote C2C12 differentiation

Myogenic differentiation of C2C12s seeded in conductive microgel scaffolds was analyzed at 3 and 7 days *via* PCR and immunocytochemistry. The early myogenic marker, MyoD (*Myod1*), did not offer a conclusive pattern in gene expression in response to microgel porosity or conductivity (**Fig. 3A**). The expression of the slightly later myogenic marker, myogenin (*Myog*), indicated cell response was unaffected by substrate conductivity, but was sensitive to hydrogel structure and time (**Fig. 3B**). While *Myog* expression was generally downregulated compared to the control, it was higher at 3 days than 7 days, as expected. No changes in gene expression were observed when cells were seeded in bulk degradable controls. Expression of the later myogenic marker, myosin heavy chain (*Myh7*), suggested potential interactions between conductivity, physical structure, and time (**Fig. 3C**). Most notably, there was a significant increase in *Mhy7* expression by cells in conductive microgel scaffolds compared to the non-conductive group at 7 days. Few differences existed between the remaining interactions, though a trend for greater *Myh7* expression was observed in the cells grown in conductive bulk gels at 7 days compared to those in non-conductive gels. When the interactions between time, physical properties, and electrical properties were analyzed with a three-way ANOVA, *Myh7* expression was influenced by the combination of time and electrical cues (*p*=0.0002) as well as time and physical cues (*p*=0.0073).

**Figure 3:**
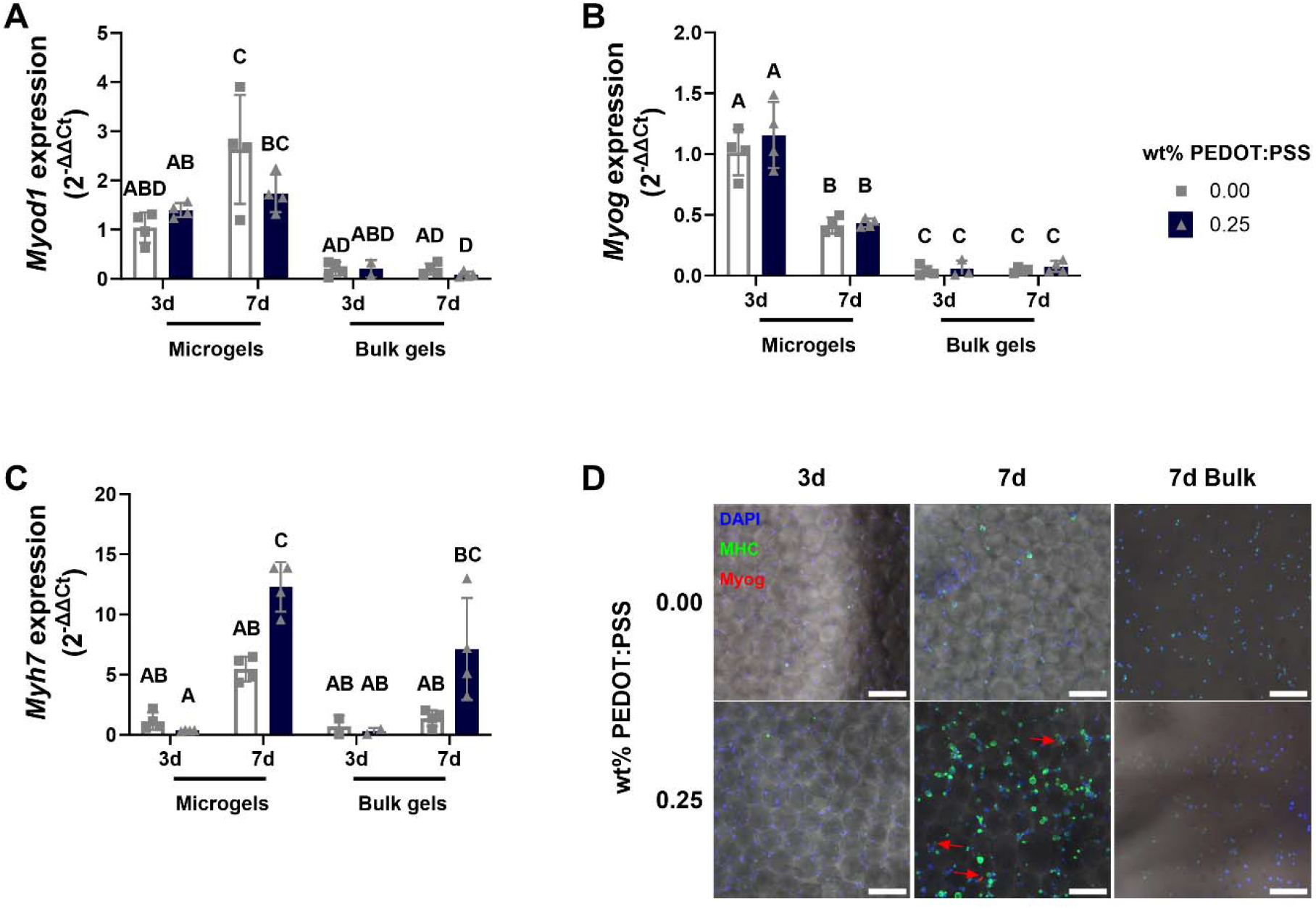
Microgel structure aids in expression of early myogenic markers of C212s, while conductivity enhances expression of late myogenic markers. **(A)** Expression of the early myogenic gene, *Myod1*, was not affected by either the electrical or physical properties of the microgel annealed scaffolds (n=3-4). **(B)** Myogenin gene expression (*Myog*) was not influenced by scaffold conductivity but was dependent on scaffold porosity and time (n=3-4). **(C)** Myosin heavy chain gene expression (*Myh7*) was influenced by conductivity, porosity, and time (n=3-4). Statistical differences were determined using a two-way ANOVA. Groups denoted with different letters are significantly different, while those that share letters are similar. **(D)** C2C12s exhibited upregulated myosin heavy chain (MHC) protein expression when in conductive microgel scaffolds. Trace myogenin (red arrows) was observed in this group but was not observable in cells grown in the non-conductive gels or bulk gels at both time points. Scale bar represents 200 µm.

Immunostaining for MHC was more pronounced in conductive microgels at 7 days compared to non-conductive controls (**Fig. 3D**). Although nuclei staining indicated good cell distribution within the microgel scaffolds, there was no discernable MHC staining at 3 days for either microgel group. C2C12s seeded in bulk degradable gels exhibited a rounded cell morphology and minimal MHC staining. Cells were also stained for myogenin, but signal was limited and only visible in conductive microgel scaffolds at 7 days. These data agree with MHC gene expression analysis and affirm that both physical and electrical properties of a material influence C2C12 differentiation.

### 3.4 Conductive microgels promote early myogenic differentiation of skMDCs

Human skeletal muscle-derived cells (skMDCs) were used as a more clinically relevant *in vitro* model to probe the role of biomaterial porosity and conductivity on myogenic differentiation. Similar to the studies performed with C2C12s, myogenic potential was interrogated on the gene and protein level. *Myod1* expression was upregulated after 1 day for cells in the conductive microgel scaffolds over the non-conductive controls (**Fig. 4A**). Since *Myod1* is involved in cell cycle arrest associated with differentiation, these data indicate differentiation is initiated earlier in conductive scaffolds. By 7 days, *Myod1* expression was higher, on average, for cells in non-conductive microgels, though the results are not significant, again suggesting differences in the timing of early differentiation. Trends indicate *Myog* expression was upregulated in cells cultured in non-conductive scaffolds (**Fig. 4B**), suggesting regulation of this myogenic gene is not responsive to conductive cues at early timepoints. Myosin heavy chain transcript levels were not detectable at 1 day, but by 7 days, trends suggest *Myh7* expression was greater by cells in the non-conductive scaffolds (**Fig. 4C**). Similar trends were observed when interrogating another isoform of myosin heavy chain, *Myh2,* which was expressed only by cells in the conductive microgel scaffolds on 1 day (**Fig. 4D**). By 7 days, *Myh2* was expressed more by cells in the non-conductive scaffolds, though trends were not statistically significant. When myosin heavy chain protein levels were assessed *via* immunostaining, we observed similar expression between the conductive and non-conductive groups at all timepoints (**Fig. 4G**). However, cells in the conductive scaffolds had fewer punctate staining patterns than those in the non-conductive group, particularly after 1 day, indicating more robust cell structures. Furthermore, there are clearer indications of multinucleated cell bodies in the conductive scaffolds at 7 days. Collectively, these observations indicate that conductive biomaterials promote expression of myogenic differentiation markers at early timepoints.

**Figure 4:**
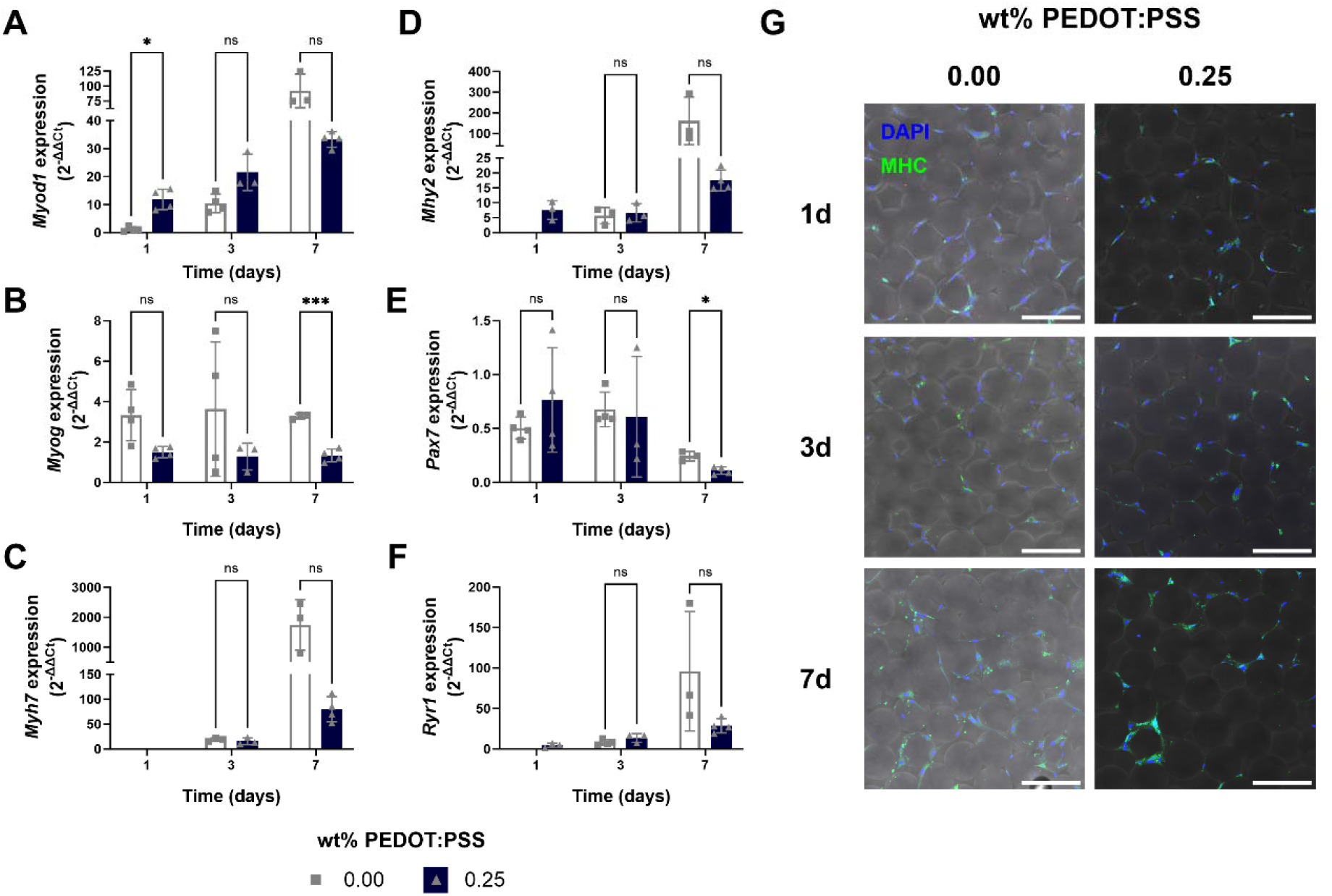
Conductive microgel scaffolds promote myogenic differentiation of skMDCs at early timepoints. **(A)** *Myod1* gene expression was significantly higher at 1 day when skMDCs were cultured in conductive microgel scaffolds, which points to enhanced initiation of myogenic differentiation (n=3-4, \**p*≤0.05). *Myod1* expression increases in the non-conductive scaffolds over 7 days. **(B)** *Myog* expression was downregulated in the conductive microgel scaffolds compared to the non-conductive group (n=3-4, \*\*\**p*≤0.001). **(C)** *Myh7* and **(D)** *Myh2* expression increased over time in both scaffolds (n=3-4). **(E)** *Pax7* expression was lower in conductive microgel scaffolds, indicating that conductivity promotes myogenic differentiation (n=3-4, \**p*≤0.05). **(F)** *Ryr1* expression increased over time for cells in both scaffolds (n=3-4). Statistical analyses of qPCR data were generated using multiple unpaired t-tests with Holm-Šídák correction where ns=not significant. **(G)** Immunofluorescence staining of skMDCs reveal similar myosin heavy chain (MHC) protein expression of cells cultured in conductive or non-conductive microgel scaffolds. Scale bar represents 200 µm.

*Pax7* is an established marker of muscle precursor cells, and reduced *Pax7* expression suggests enhanced myogenic differentiation. The significantly reduced expression of *Pax7* by cells in conductive scaffolds at 7 days corroborates these results (**Fig. 4E**). We also interrogated the expression of ryanodine receptor 1 (*Ryr1*), given previous literature demonstrating links between conductive biomaterials and ion channel regulation.^[24]^ Ryanodine receptors are involved in the release of calcium ions during skeletal muscle contraction, and while there were no significant differences between groups, *Ryr1* was only expressed by cells in the conductive microgel scaffolds at 1 day (**Fig. 4F**). *Ryr1* expression in conductive microgel scaffolds was more than 60% greater (2^-ΔΔCt^ = 13.62 ± 5.44, n= 3) than in non-conductive scaffolds (2^-ΔΔCt^ = 8.22 ± 3.43, n= 4). Therefore, the results of **Figure 4** suggest conductive microgel scaffolds promote myogenic differentiation of a clinically relevant cell model at early timepoints.

### 3.5 Conductive microgel scaffolds enhance secretion of cytokines related to myogenic differentiation and wound healing

Myogenic gene and protein expression differed slightly in skMDCs compared to the clear correlation between gene and protein expression in C2C12s. Therefore, we investigated if the skMDCs secreted other myogenic or regenerative factors in response to conductivity. After investigating 80 different analytes *via* a multiplex protein array, we identified six factors expressed by skMDCs in greater quantities in conductive microgel scaffolds during at least at one time point. IL-6 (**Fig. 5A**) and IL-8 (**Fig. 5B**) are pro-inflammatory cytokines that are critical for proper wound healing. Both factors were secreted in increased quantities by cells in conductive scaffolds at 3 days, while at the other time points, these factors were higher in the non-conductive scaffolds. Pro-epidermal growth factor (EGF) was expressed similarly by cells in conductive and non-conductive scaffolds, except at 3 days where there was a 1.5-fold increase between groups (**Fig. 5C**). Muscle cells in conductive scaffolds secreted more than twice as much VEGF at day 1, with similar levels secreted at 3 and 7 days (**Fig. 5D**). IGF-1 is implicated in muscle cell growth *via* hypertrophy, and IGFBP-2 prolongs the half-life of IGF-1. Insulin-like growth factor (IGF-1) was expressed more by cells in conductive scaffolds at 1 day, but the increase was even more prominent at 3 days (**Fig. 5E**). IGF-binding protein (IGFBP-2) was also expressed more by cells in conductive microgel scaffolds on 1 and 7 days (**Fig. 5F**). These data indicate that skMDCs have greater myogenic potential at early time points when cultured in conductive scaffolds.

**Figure 5:**
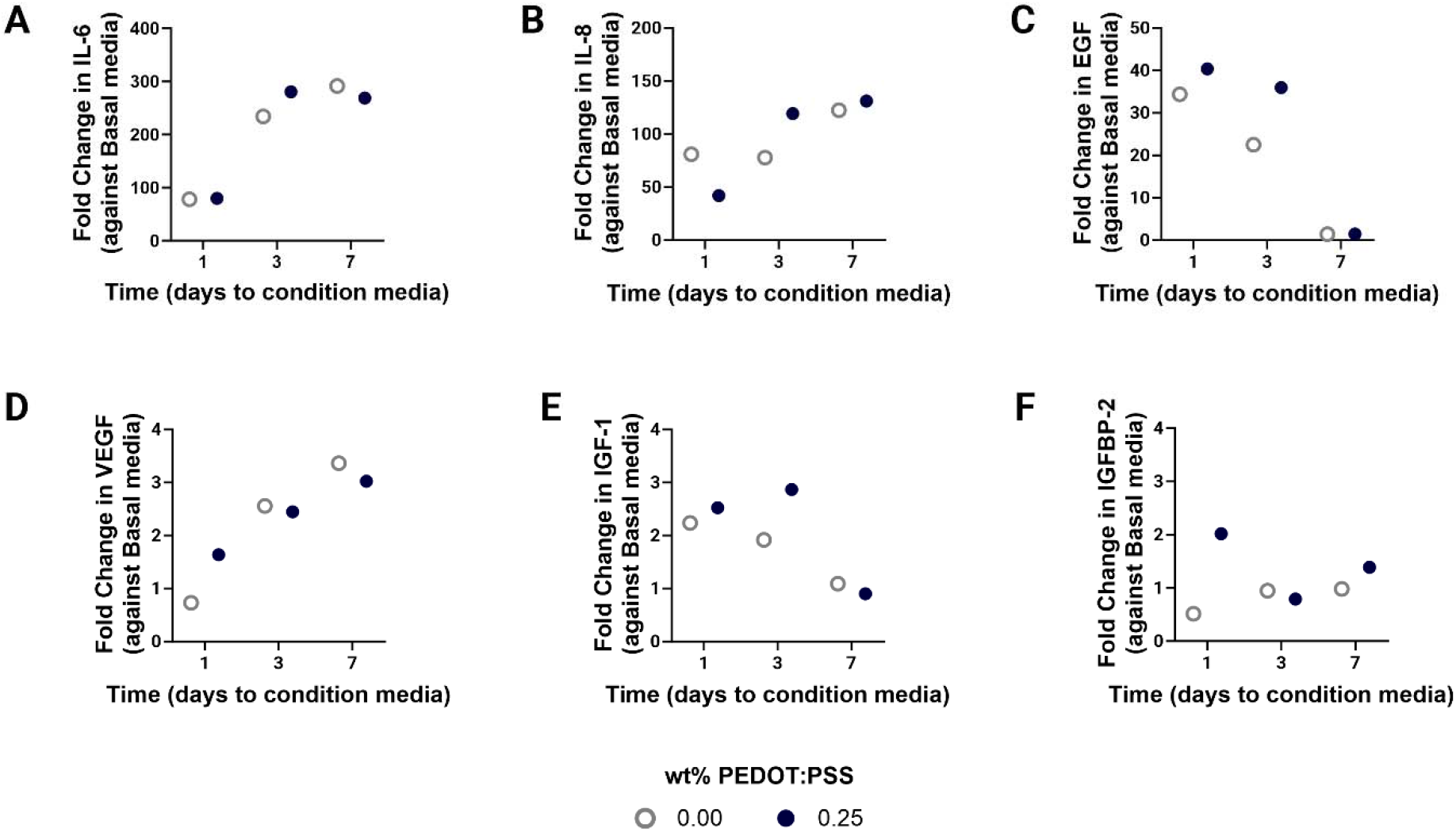
Conductive microgel scaffolds enhance early secretion of cytokines related to myogenic differentiation and wound healing. **(A)** IL-6 and **(B)** IL-8 were secreted more by skMDCs in the conductive microgel scaffolds at 3 days. skMDCs secreted more **(C)** EGF and **(D)** VEGF in conductive microgels at early timepoints. skMDCs secreted **(E)** IGF-1 and **(F)** IGFBP-2 at least 1.5-fold more when cultured in conductive microporous scaffolds at early timepoints, which has positive implications for early muscle cell repair. Statistical comparison is not possible due to singular replicates for each analyte.

### 3.6 Late-stage factors secreted by skMDCs reduced M1 macrophage polarization

Macrophages play a pivotal role in muscle homeostasis, injury, and repair,^[25]^ so we assessed the functional effects of skMDC-secreted factors on macrophages. We treated human THP-1 macrophages with media conditioned by skMDCs cultured in conductive microgel scaffolds for 1, 3, and 7 days, and analyzed macrophage polarization *via* flow cytometry. We found high macrophage viability (>70%) for both conductive and nonconductive groups across all timepoints (**Fig. 6A**). We also observed a subtle decrease in overall M1 polarized macrophages over time, where media from skMDCs cultured on nonconductive microgels resulted in slightly increased M1 polarization compared to those cultured on conductive gels (**Fig. 6B**). In contrast, M2 macrophages were relatively low for both scaffolds at 1 day but nearly doubled when treated with 3 day-conditioned media. Interestingly for 7-day groups, we noted significantly decreased M2 macrophages only for those treated with conditioned media from skMDCs cultured on conductive microgels (**Fig. 6C**). These M2 macrophage trends correspond with M2-associated cytokines, such as IL-4, IL-10, and G-CSF, identified in the protein array (**Fig. S2**). Additionally, this correlates with IL-6 secretion (**Fig. 5A**), which, as a pleiotropic cytokine, can produce an anti-inflammatory effect under certain conditions.^[26]^ Together, these data reveal a dynamic relationship between myogenic cells and the immune microenvironment that is dependent on time and material conductivity.

**Figure 6.**
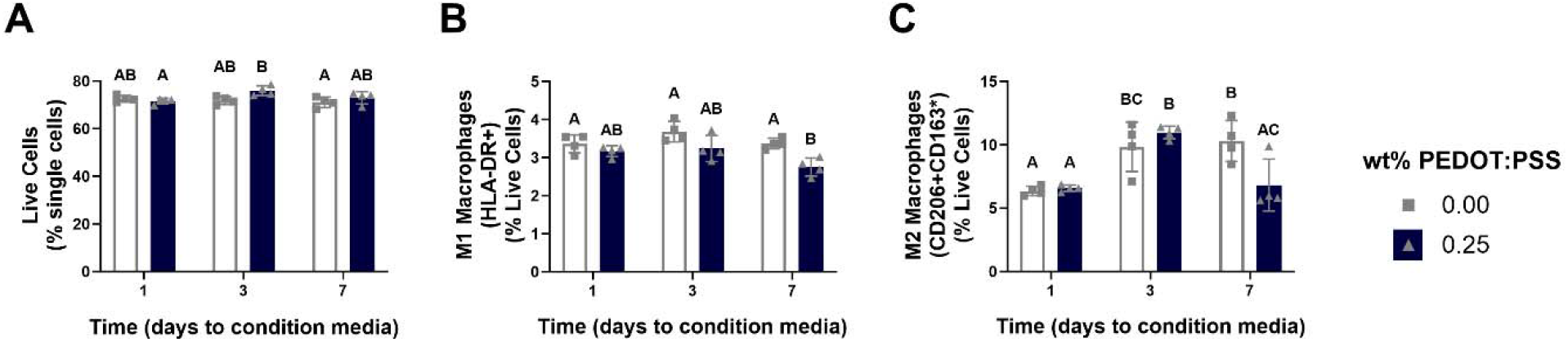
Late-stage skMDC-secreted factors reduced M1 macrophage polarization. Flow cytometric analyses of human THP-1 macrophages treated with skMDC conditioned media for 24 hours. Quantifications of **(A)** live cells, **(B)** HLA-DR^+^ (M1) macrophages, and **(C)** CD206^+^CD163^+^ (M2) macrophages. Groups with statistically significant differences based on two-way ANOVA do not share the same letters.

## 4. Discussion

Biophysical and electrical properties of biomaterials used as cell carriers are critical for directing cell behavior and can synergistically promote cell differentiation and maturation. In this work, we developed conductive microgel annealed scaffolds to interrogate myogenic differentiation of murine and human myoblastic cells. Unlike traditional bulk hydrogels, microgel annealed scaffolds possess inherent void space between the particles which permit immediate cell migration. Previously, gels formed entirely of PEDOT:PSS were fragmented into injectable microgels, and produced a limited inflammatory response when implanted *in vivo*.^[27]^ Conductive microgels have also been formed by gallol modification of hyaluronic acid and induced muscle contractility *ex vivo*.^[7]^ However, neither of these studies interrogated the influence of conductive microgels on myoblast differentiation. Herein, we demonstrate the ability of conductive microgel annealed scaffolds with dispersed PEDOT:PSS to enhance myoblast differentiation and maturity. The monodisperse nature of microfluidic-produced microgels, as employed here, ensures consistent, predictable, and tunable void space and resultant material properties.

PEDOT:PSS is frequently used for fabricating conductive biomaterials owing to its commercial availability and its dispersant nature when suspended in water. The hydrophobic PEDOT^+^ core is surrounded by a shell of PSS^-^, which forms micelles that can evenly distribute within water-based materials such as hydrogels.^[28]^ This contrasts starkly to other commonly used synthetic conductive materials such as polypyrrole, polyaniline, and graphene. Despite high levels of electrical conductivity, these materials require further chemical processing to overcome their hydrophobic properties for incorporation into hydrogels. Although addition of PEDOT:PSS into the PEG microgels doubled the conductivity of annealed scaffolds, compressive moduli of individual microgels or the storage modulus of annealed scaffolds was unchanged.^[18]^ Thus, the incorporation of PEDOT:PSS allowed us to successfully decouple electrical and mechanical properties and facilitate the interrogation of how these properties individually influence cell behavior.

Bioelectrical components have been previously incorporated into hydrogels for use in muscle tissue engineering, which achieved similar electrical properties as reported here.^[11]^ The PEDOT:PSS-containing microgel scaffold conductivity was approximately 3.5 x 10^-6^ S/cm, which is of similar magnitude with other studies using polypyrrole and collagen (1.5 x 10^-5^ S/cm),^[15]^ PEDOT:PSS (6.1 x 10^-6^ S/cm),^[29]^ and aniline-based polymers with chitosan (3.5 x 10^-5^ S/cm).^[30]^ Lower conductivities reported in this study may be due to how PEDOT:PSS was dispersed within the PEG microgels. This dispersion could result in greater π-π orbital distances, thereby limiting free electron transfer.^[7]^ Future studies could examine how decorating a material’s surface with conductive moieties may promote enhanced electrical interactions at the cell-material interface. Additionally, though each of these studies characterized changes in mechanical properties with the inclusion of conductive additives, few specifically interrogate the interplay of biomaterial electrical and physical properties on influencing cell response. The novelty of this work includes examining microporosity as a physical property and how it interacts with electroactivity to affect cell behavior and myogenic differentiation. While reports extolling the benefits of conductive biomaterials on cell behavior are increasing, there are currently no standardizations for measuring the electroactive properties of biomaterials. This hinders accurate comparison between materials across studies and remains a significant knowledge gap in the field of electroactive materials.

The results of these studies established that both microporosity and conductivity were essential for directing C2C12 and human skMDC myoblast differentiation. Microgel annealed scaffolds promoted increased proliferation and gene expression compared to bulk scaffolds. When probing for myogenic markers, we observed a stark increase in myosin heavy chain gene and protein expression in C2C12s grown in conductive scaffolds at later time points. Myosin heavy chain is a hallmark indicator for muscle cells that are maturing from myoblasts to more functional myotubes. In skMDCs, *Myh7* was not differentially expressed, though this may be due to the prevalence of *Myh7* in cardiomyocyte function, rather than skeletal muscle cells. As such, we also tested how skMDCs expressed *Myh2*. At day 1, only those cells cultured in conductive microgel scaffolds expressed *Myh2*, suggesting the ability of conductive biomaterials to enhance myogenesis potential at early timepoints. Furthermore, reduced *Pax7* expression at 7 days suggests conductive microporous scaffolds improve myogenic differentiation at later timepoints. Early stage markers of myogenic differentiation (*e.g.*, MyoD and myogenin) were not as dependent on conductivity as later-stage markers.^[31–33]^] However, since our hypothesis was that conductive microporous materials would promote myogenic differentiation, it is possible that these events occurred at earlier time points.

Conductive biomaterials direct bioelectric signaling within cells and tissues through many avenues including regulating ion channels in the cell membrane.^[8]^ As such, we tested the expression of ryanodine receptor 1, which is involved in the release of calcium ions during cardiac and skeletal muscle cell contraction.^[24]^ Only cells cultured in conductive microgel scaffolds expressed *Ryr1* on day 1, and although not significant, there was a nearly 2-fold increase in *Ryr1* expression by cells grown in conductive scaffolds over non-conductive controls at 3 days. These results indicate that improvements in myogenesis on conductive materials may be, in part, explained by changes in ion channel regulation in response to material electrical cues.

We further investigated changes in the skMDC secretome to determine whether conductivity influences secreted myogenic or regenerative factors. IGF-1 and IGFBP-2 are growth factors associated with myofiber hypertrophy and regeneration in skeletal muscle,^[34–37]^] and both were secreted more by cells in conductive microgel scaffolds at early time points. VEGF expression was also upregulated early when cells were cultured in conductive microgel scaffolds. We validated VEGF bioactivity by assessing tubule formation by cells treated with skMDC-conditioned media. Tubulogenesis was similar when endothelial cells were treated with media conditioned by cells in conductive and non-conductive materials, though trends suggested enhanced branching in response to factors secreted by cells in conductive scaffolds at early time points. These trends were also expected, given that differences in the regenerative behavior of skMDCs in response to conductive cues are subtle in this work.

Macrophage-mediated muscle regeneration is a well characterized process, and our data largely correlate with trends observed *in vivo*.^[25,38]^ When THP-1 macrophages were treated with conditioned media from skMDCs cultured on conductive microgels, we observed initial high levels of M1 phenotypes that decreased over time, as well as early low levels of M2 macrophages that peaked when treated with 3 day-conditioned media, then tapered at 7 days. This is representative of a muscle wound response, which is characterized by high levels of early inflammation that are replaced over 1-2 weeks by pro-regenerative mediators.^[25,39]^ Furthermore, the decrease in M2 polarized macrophages when treated with 7 day-conditioned media could indicate advanced resolution of this response mechanism and decreased likelihood of a fibrotic response, which is often associated with the persistence of M2-related cytokines, such as IL-10.^[40]^ Given the serial nature of these experiments, these data indicate that cytokine secretions from skMDCs cultured on conductive microgels not only encourage immune-mediated muscle repair mechanisms, but they may in fact direct them. Future work is warranted using *in vivo* models of muscle repair to explore the translation of this finding to a wound site.

While the microporosity of these gels provides exciting opportunities to enhance cell proliferation and migration, microgel annealed scaffolds are limited by the random porosity inherent in their structure. Surface topography and muscle cell alignment is vital in myogenic differentiation due to the highly organized structure of muscle.^[3,41]^ Future work may consider orienting the microgels into an aligned structure through the application of an external electric field^[42–44]^] or bioprinting.^[45–47]^] Additionally, while MHC expression was visibly upregulated on conductive microgels, we did not observe myocyte fusion into myotubes. This may be improved with higher seeding density into the annealed constructs or incorporating mechanisms for the material to degrade and thus make room for myotubes to self-assemble.

To our knowledge, this is the first report that decouples electrical conductivity and microporosity and investigates the interplay of these properties on cellular myogenic response. Both electrical and biophysical properties were integral for promoting myoblast differentiation at both the gene and protein level. The bioactivity and injectable nature of the microgel scaffolds make them a promising tool for clinical translation to heal muscle wounds. Future work will investigate the translation of this platform in an *in* vivo model such as volumetric muscle loss.

## Acknowledgements

This work was supported by the National Institutes of Health (R01 AR079211) to JKL. JKL gratefully acknowledges financial support from the Lawrence J. Ellison Endowed Chair of Musculoskeletal Research. AC was funded by the Floyd and Mary Schwall Fellowship in Medical Research, the University of California Davis Dean’s Distinguished Graduate Fellowship, and the ARCS Foundation. KHG was funded by the Maxine Adler Graduate Fellowship from the Graduate Student Support Program at the University of California, Davis, School of Veterinary Medicine. ACF was supported by a National Institute of Arthritis and Musculoskeletal and Skin Diseases funded training program in Musculoskeletal Health Research (T32 AR079099). We gratefully acknowledge the contributions of Dr. David Ramos-Rodriguez for technical assistance. Part of this study was carried out at the UC Davis Center for Nano and Micro Manufacturing (CNM2).

## Data availability statement

Data available upon request from the authors.

## SUPPLEMENTAL MATERIALS FOR

**Figure S1.**
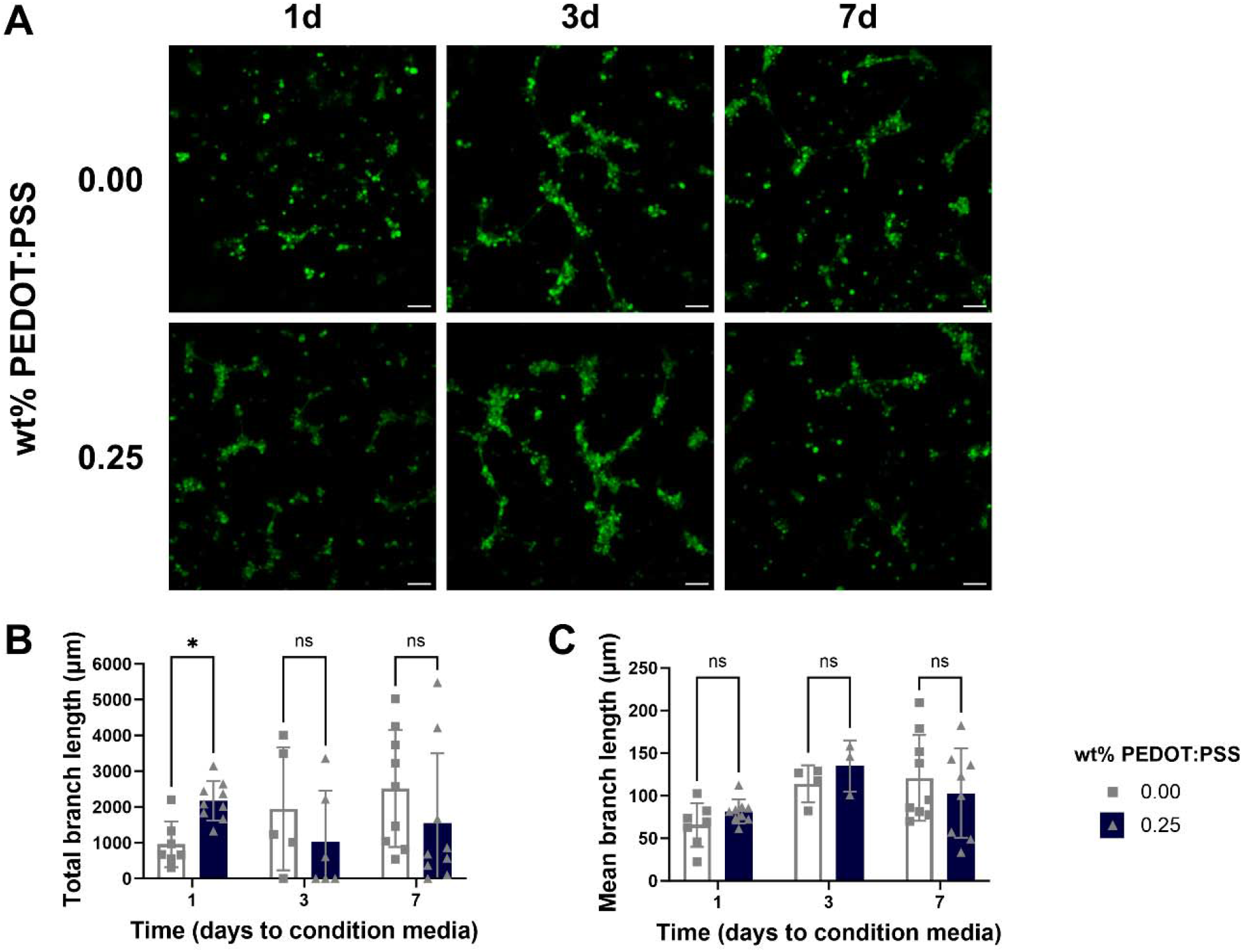
Early muscle cell-secreted factors promote endothelial cell tube formation. **(A)** Representative images of tubules after 6 hours (scale = 100 µm). HMVECs exhibited more branch formation with media conditioned by skMDCs in conductive microgel scaffolds for 1 day compared to media from non-conductive scaffolds. More robust branching was observed when HMVECs were treated with 3 day-conditioned media, regardless of conductivity. **(B)** Total branch length was higher for HMVECs treated with media conditioned by skMDCs on conductive microgel scaffolds for 1 day, corroborating observations in **(A)**. No differences were observed when total branch length was analyzed as a function of conductivity at the other durations of media conditioning (n=5-9). **(C)** Average branch length was similar between groups at each time point (n=3-9). Groups were compared using multiple, unpaired t-tests where \**p*≤0.05 and ns = not significant.

**Figure S2:**
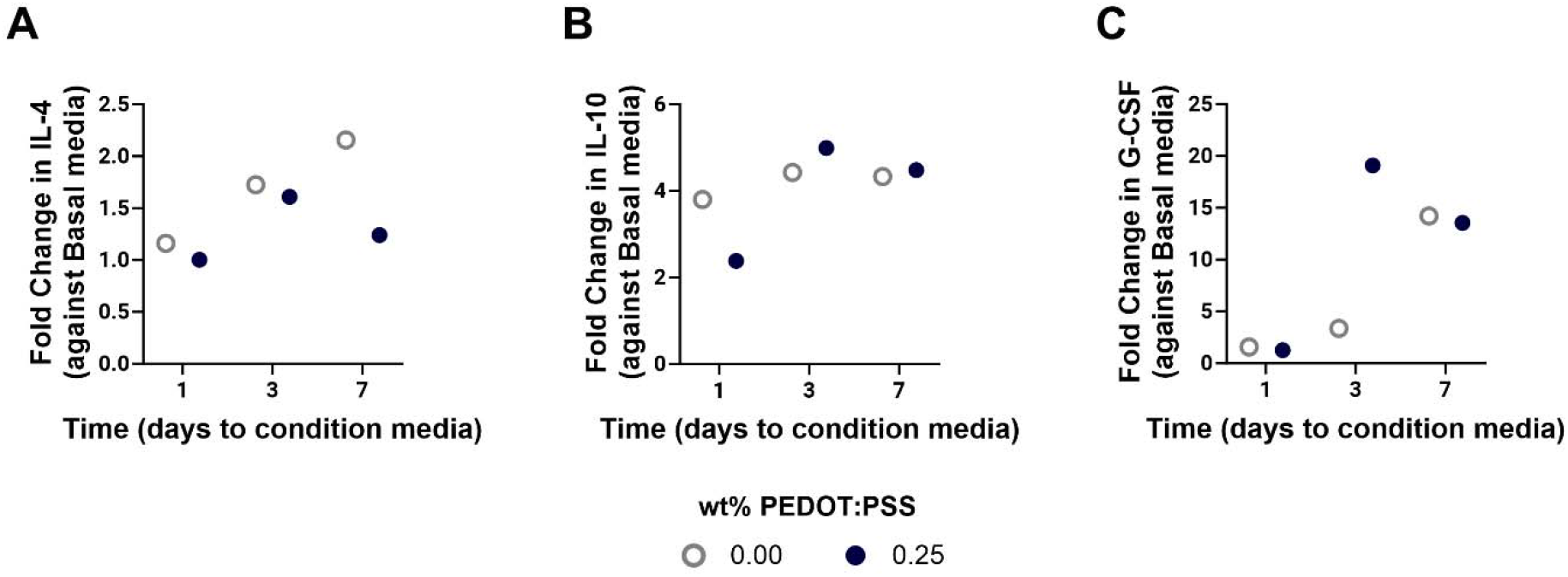
IL-4, IL-10, and G-CSF secretion from skMDCs cultured in microgels for 1, 3, and 7 days. Relative secretion of **(A)** IL-4, **(B)** IL-10, and **(C)** G-CSF as evaluated *via* protein array. Statistical comparison is not possible due to singular replicates for each analyte.

## References

[1] B. T. Corona, J. C. Rivera, J. G. Owens, J. C. Wenke, C. R. Rathbone, J Rehabil Res Dev 2015, 52, 785.

[2] H. Manring, E. Abreu, L. Brotto, N. Weisleder, M. Brotto, Front. Physiol. 2014, 5.

[3] S. Jana, S. K. L. Levengood, M. Zhang, Adv. Mater. 2016, 28, 10588.

[4] J. M. Grasman, M. J. Zayas, R. L. Page, G. D. Pins, Acta Biomater. 2015, 25, 2.

[5] R. Lev, D. Seliktar, *J. R. Soc.* Interface 2018, 15, 20170380.

[6] J. P. Newsom, K. A. Payne, M. D. Krebs, Acta Biomater. 2019, 88, 32.

[7] M. Shin, K. H. Song, J. C. Burrell, D. K. Cullen, J. A. Burdick, Adv. Sci. 2019, 6, 1901229.

[8] A. Casella, A. Panitch, J. K. Leach, Bioelectricity 2021, 3, 27.

[9] X. Liu, A. L. M. Ii, S. Park, B. E. Waletzki, A. Terzic, M. J. Yaszemski, L. Lu, *J. Mater. Chem.* B 2016, 4, 6930.

[10] B. W. Walker, R. P. Lara, C. H. Yu, E. S. Sani, W. Kimball, S. Joyce, N. Annabi, Biomaterials 2019, 207, 89.

[11] R. Dong, P. X. Ma, B. Guo, Biomaterials 2020, 229, 119584.

[12] M.-C. Chen, Y.-C. Sun, Y.-H. Chen, Acta Biomater. 2013, 9, 5562.

[13] S. Hosseinzadeh, M. Mahmoudifard, F. Mohamadyar-Toupkanlou, M. Dodel, A. Hajarizadeh, M. Adabi, M. Soleimani, Bioprocess Biosyst. Eng. 2016, 39, 1163.

[14] K. D. McKeon-Fischer, D. P. Browe, R. M. Olabisi, J. W. Freeman, *J. Biomed. Mater. Res.* A 2015, 103, 3633.

[15] I. M. Basurto, M. T. Mora, G. M. Gardner, G. J. Christ, S. R. Caliari, Biomater. Sci. 2021, 9, 4040.

[16] J. M. de Rutte, J. Koh, D. Di Carlo, Adv. Funct. Mater. 2019, 29, 1900071.

[17] J. M. Lowen, G. C. Bond, K. H. Griffin, N. K. Shimamoto, V. L. Thai, J. K. Leach, Adv. Healthc. Mater. 2023, 2202239.

[18] A. Casella, A. Panitch, J. K. Leach, *J. Biomed. Mater. Res.* A 2023, 111, 596.

[19] T. Gonzalez-Fernandez, A. J. Tenorio, A. M. Saiz Jr, J. K. Leach, Adv. Healthc. Mater. 2022, 11, 2102337.

[20] G. Carpentier, *Protein Array Analyzer for ImageJ*, 2010.

[21] G. Carpentier, S. Berndt, S. Ferratge, W. Rasband, M. Cuendet, G. Uzan, P. Albanese, Sci. Rep. 2020, 10, 11568.

[22] P. H. Mott, C. M. Roland, *Phys. Rev.* B 2009, 80, 132104.

[23] T. Gonzalez-Fernandez, P. Sikorski, J. K. Leach, Acta Biomater. 2019, 96, 20.

[24] J. T. Lanner, D. K. Georgiou, A. D. Joshi, S. L. Hamilton, Cold Spring Harb. Perspect. Biol. 2010, 2, a003996.

[25] X. Wang, L. Zhou, Front. Cell Dev. Biol. 2022, 10, 952249.

[26] J. Mauer, B. Chaurasia, J. Goldau, M. C. Vogt, J. Ruud, K. D. Nguyen, S. Theurich, A. C. Hausen, J. Schmitz, H. S. Brönneke, E. Estevez, T. L. Allen, A. Mesaros, L. Partridge, M. A. Febbraio, A. Chawla, F. T. Wunderlich, J. C. Brüning, Nat. Immunol. 2014, 15, 423.

[27] V. R. Feig, S. Santhanam, K. W. McConnell, K. Liu, M. Azadian, L. G. Brunel, Z. Huang, H. Tran, P. M. George, Z. Bao, Adv. Mater. Technol. 2021, 6, 2100162.

[28] S. Zhang, Y. Chen, H. Liu, Z. Wang, H. Ling, C. Wang, J. Ni, B. Celebi-Saltik, X. Wang, X. Meng, H. J. Kim, A. Baidya, S. Ahadian, N. Ashammakhi, M. R. Dokmeci, J. Travas-Sejdic, A. Khademhosseini, Adv Mater 2020, 32, e1904752.

[29] A. G. Guex, J. L. Puetzer, A. Armgarth, E. Littmann, E. Stavrinidou, E. P. Giannelis, G. G. Malliaras, M. M. Stevens, Acta Biomater. 2017, 62, 91.

[30] B. Guo, J. Qu, X. Zhao, M. Zhang, Acta Biomater. 2019, 84, 180.

[31] H. Hwangbo, H. Lee, E. Jin, Y. Jo, J. Son, H. M. Woo, D. Ryu, G. H. Kim, Adv. Funct. Mater. 2023, 33, 2209157.

[32] Y.-N. Jang, E. J. Baik, JAK-STAT 2013, 2, e23282.

[33] P. Fortini, E. Iorio, E. Dogliotti, C. Isidoro, Front. Physiol. 2016, 7, 237.

[34] M. E. Coleman, F. DeMayo, K. C. Yin, H. M. Lee, R. Geske, C. Montgomery, R. J. Schwartz, J. Biol. Chem. 1995, 270, 12109.

[35] A. Musarò, K. McCullagh, A. Paul, L. Houghton, G. Dobrowolny, M. Molinaro, E. R. Barton, H. L Sweeney, N. Rosenthal, Nat. Genet. 2001, 27, 195.

[36] A. P. Sharples, N. Al-Shanti, D. C. Hughes, M. P. Lewis, C. E. Stewart, Growth Horm. IGF Res. 2013, 23, 53.

[37] C. W. Ernst, D. C. McFarland, M. E. White, Differentiation 1996, 61, 25.

[38] M. Shang, F. Cappellesso, R. Amorim, J. Serneels, F. Virga, G. Eelen, S. Carobbio, M. Y. Rincon, P. Maechler, K. De Bock, P.-C. Ho, M. Sandri, B. Ghesquière, P. Carmeliet, M. Di Matteo, E. Berardi, M. Mazzone, Nature 2020, 587, 626.

[39] R. Furrer, P. S. Eisele, A. Schmidt, M. Beer, C. Handschin, Sci. Rep. 2017, 7, 40789.

[40] C. Cui, R. K. Driscoll, Y. Piao, C. W. Chia, M. Gorospe, L. Ferrucci, Aging Cell 2019, 18.

[41] H. Gao, J. Xiao, Y. Wei, H. Wang, H. Wan, S. Liu, ACS Omega 2021, *6*, 20931.

[42] W. Kim, H. Lee, C. K. Lee, J. W. Kyung, S. B. An, I.-B. Han, G. H. Kim, Adv. Funct. Mater. 2021, 31, 2105170.

[43] I.-C. Liao, J. B. Liu, N. Bursac, K. W. Leong, Cell. Mol. Bioeng. 2008, 1, 133.

[44] M. Flaibani, L. Boldrin, E. Cimetta, M. Piccoli, P. D. Coppi, N. Elvassore, *Tissue Eng.* Part A 2009, 15, 2447.

[45] S. Ostrovidov, S. Salehi, M. Costantini, K. Suthiwanich, M. Ebrahimi, R. B. Sadeghian, T. Fujie, X. Shi, S. Cannata, C. Gargioli, A. Tamayol, M. R. Dokmeci, G. Orive, W. Swieszkowski, A. Khademhosseini, Small 2019, 15, 1805530.

[46] T. Fan, S. Wang, Z. Jiang, S. Ji, W. Cao, W. Liu, Y. Ji, Y. Li, N. Shyh-Chang, Q. Gu, Biofabrication 2021, 14, 015009.

[47] G. H. Yang, W. Kim, J. Kim, G. Kim, Theranostics 2021, 11, 48.

